# Scaling laws of human transcriptional activity

**DOI:** 10.1101/2023.08.10.551625

**Authors:** Jiayin Hong, Ayse Derya Cavga, Devina Shah, Ernest Laue, Jussi Taipale

## Abstract

Each human chromosome maintains its individuality during the cell cycle, and occupies a spatially limited volume, termed chromosome territory. Each linear chromosomal DNA is folded into multiple loops in the three dimensional space, and further organized into densely packed heterochromatin, less dense euchromatin and nucleosome-free regions that are accessible for transcription factor binding. As the average density of chromatin in the nucleus is very high, size exclusion potentially restricts access of large macromolecules such as RNA polymerase II and Mediator to DNA buried in chromosomal interiors. To examine this idea, we investigated whether increase in chromosome size leads to relative decrease in transcriptional activity of larger chromosomes. We found that the scaling of gene expression relative to chromosome size follows exactly the surface-area-to-volume ratio, suggesting that active genes are located at chromosomal surfaces. To directly test this hypothesis, we developed a scalable probe to assess chromatin accessibility to macromolecules of different sizes. We show that, at the chromosomal level, open chromatin landscapes of small and large molecules are strikingly similar. However, at a finer locus level, regions accessible to small transcription factors were primarily enriched around promoters, whereas regions accessible to large molecules were dispersed along gene bodies. Collectively, our results indicate that DNA accessibility is controlled at two different scales, and suggest that making chromatin accessible to large molecules is a critical step in the control of gene expression.

## Introduction

Proper regulation of gene expression is critical for cellular processes such as proliferation and differentiation, and abnormal gene expression can lead to disease. It is well established that chromatin architecture plays a crucial role in gene regulation^1–3^. Despite considerable progress that has been made in understanding the 3D organization of chromosomes^4,5^, and how it contributes to control of gene expression^6^, there remains a gap in our knowledge regarding chromatin accessibility to large macromolecules.

Macromolecules involved in transcription, such as transcription factors (TFs), exhibit significant variability in size. TFs, with diameters of approximately 3-5 nm, can efficiently navigate the genome and bind to regulatory elements even within condensed chromatin domains. In contrast, it has been suggested that larger molecular complexes such as RNA polymerase II (Pol II) and Mediator, measuring around 20 nm in diameter, have difficulty to access densely packed chromatin domains. This physical constraint imposed on transcription is known as size exclusion effect^7^ (**Fig. 1a**).

**Fig. 1:**
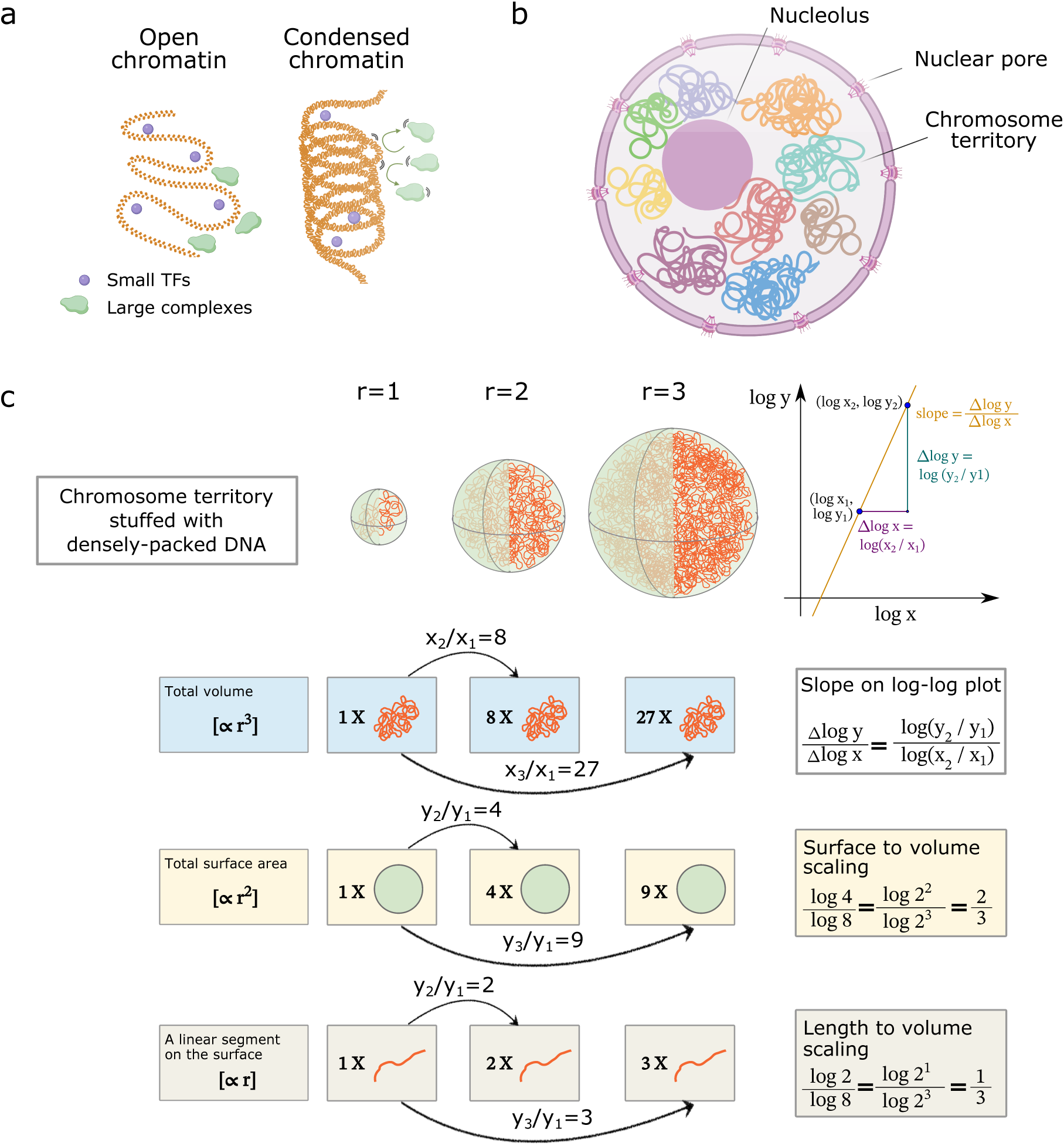
Size exclusion effect and territorial chromosome lead to the sublinear scaling of transcriptional activity relative to chromosome size. **a,** Illustration of size exclusion effect. Large complexes are likely to be hindered at the surface of condensed chromatin to access densely packed DNA but they are able to access open chromatin regions, whereas small TFs can more easily diffuse into condensed chromatin domains. **b,** Diagram of chromosome territory in the interphase nucleus. Chromatin is folded into multiple loops and each chromosome occupies a distinct territory. **c,** The scaling of volume, surface area and linear segment as chromosome size increases. The shape of a chromosome territory is approximated by a sphere. When the radius of the sphere varies from 1, 2, to 3 units, the total volume that the sphere encloses is proportional to the linear dimension cubed, whereas the total surface area is proportional to the linear dimension squared and a segment that is confined in a linear dimension just varies linearly. As a result, the slope of the straight line capturing such power-law relationships on a log-log plot would be ⅔ in the surface to volume scaling, and ⅓ in the length to volume scaling.

However, the widely used assays for chromatin accessibility at present (e.g., ATAC-seq, DNase-seq, MNase-seq) interrogate open chromatin regions based on the accessibility to transposase, DNase I, micrococcal nuclease, etc^8–10^, all of which are relatively small proteins. For instance, Tn5 transposase, a commonly used probe in ATAC-seq, is a 53.3 kDa protein^11^ with an estimated diameter of 6.8 nm^12^. Consequently, these assays inherently overlook the increased steric hindrance experienced by larger biomolecules. To address this limitation and gain insights into the regions accessible to larger molecules, as well as to study the impact of size exclusion on transcriptional regulation, we have developed a scalable probe for assessing chromatin accessibility. This novel probe mimics the behavior of molecules of different sizes scanning and searching the human genome.

## Results

### The sublinear scaling of human transcriptional activity with respect to chromosome size

Chromosomes in interphase exist as relatively compact structures and their arrangement is confined by the boundaries of the cell nucleus. Each chromosome maintains its individuality during the cell cycle and occupies a spatially limited volume; this feature of nuclear architecture is known as chromosome territory^13^ (**Fig. 1b**). It has been proposed that the outer surface of an individual chromosome territory generally provides a particular compartment for gene dense and/or transcriptionally active domains^14^. We therefore hypothesized that large macromolecular complexes like Pol II and Mediator cannot enter condensed chromatin domains in the interior of the chromosome territories. From this it would follow that transcription is preferentially located at the accessible surface of the territories. The hypothesis is testable, as it predicts that smaller chromosomes would have more gene expression per unit size.

As a chromosome increases in diameter, the volume of its territory increases cubically, hence the genetic material it can contain also increases cubically. In contrast, the amount of genetic material accessible on the surface of chromosomes increases proportionately less than the volume, because surface area is proportional to the square of diameter. If only the chromosome surface would be accessible to large macromolecules, increase in chromosome size would lead to a lower fraction of the chromosome being accessible to the transcription machinery. Theoretically, the scaling between gene expression and chromosome size would follow a power law, where the fold change of gene expression would be proportional to the fold change of chromosome size raised to a fixed exponent. The signature of a power law relationship is a straight line joining logarithmically transformed variables, whose slope is the exponent of the power law. In the case of surface to volume scaling, the slope would be equal to ⅔, the surface-area-to-volume ratio. Finally, as a chromosome increases in size, a segment that is confined in a linear dimension (e.g., genes being expressed show increased stiffness and rigidity due to decoration with bulky nascent ribonucleoproteins^15^) would increase linearly in proportion with the diameter of the chromosome territory. In other words, the exponent of the power law would be equal to ⅓ for a length to volume scaling (**Fig. 1c**).

Compared to a linear scaling relationship where both quantities increase or decrease to the same extent, in which case the slope would be exactly 1, the less-than-one scaling relationships, i.e. sublinear scaling, indicate that the bigger the chromosomes are, the less transcriptional activity they would have per length of DNA packaged into them. Therefore, a smaller chromosome would have a greater ratio of surface area to volume, and thus be less susceptible to the size exclusion effect.

To test our assumption of sublinear scaling law of transcriptional activity, we first examined the relationship between Transcription Start Sites (TSSs) and chromosome size. We used the count of TSSs on each chromosome as a proxy of transcriptional activity, and plotted the logarithmically transformed counts of TSSs against the logarithmically transformed chromosome size in base pair (**Fig. 2a**). Noticeably, the slope of the regression line is 0.67, which is quite close to the ⅔ surface-area-to-volume ratio. The observed correlation was surprisingly high, given that it is well established that many other variables, such as GC content^16^ and presence of repeats affect gene density within individual chromosomes.

**Fig. 2:**
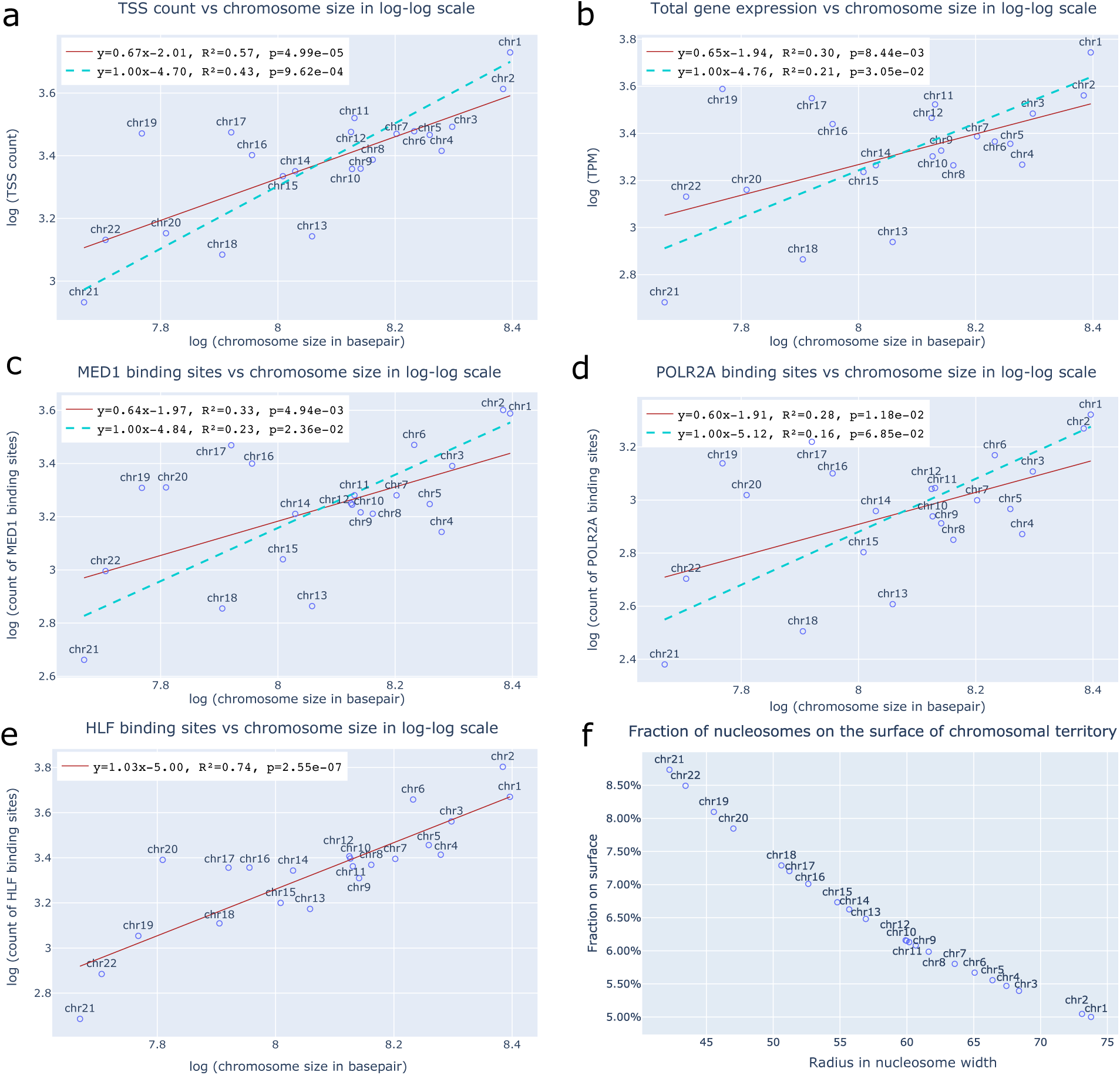
Scaling laws of transcriptional activity, TF and large complex binding, and fraction of nucleosomes on the surface. **a,** TSS count scales sublinearly to chromosome size with an exponent close to ⅔. Red solid line denotes the regression line between TSS count and chromosome size in logarithmic scale. Cyan dashed line denotes the constrained linear regression line (slope is constrained to be 1 while intercept is optimized to best fit the data, same for **b, c,** and **d**). **b,** Total gene expression scales sublinearly to chromosome size with an exponent close to ⅔. Total gene expression was quantified by TPM. **c,** The count of MED1 binding sites scales sublinearly to chromosome size with an exponent close to ⅔. The count of binding sites was determined from ChIP-seq data (same for **d** and **e**). **d,** The count of POLR2A binding sites scales sublinearly to chromosome size with an exponent close to ⅔. **e,** The count of HLF binding sites scales linearly to chromosome size with an exponent close to 1. **f,** Fraction of nucleosomes on the surface of chromosome territory as radius of the territory varies. The chromosome territory was approximated by a sphere packed with smaller spheres - nucleosomes.

To distinguish sublinear and linear relationships between count of TSSs and chromosome size, we calculated Bayesian Information Criterion (BIC) of the two models. BIC is based on the likelihood function and used as a criterion for model selection. Models with lower BIC are generally preferred. While the sublinear regression had a BIC of -25.05, the linear regression model had a BIC of -18.81. With a ΔBIC = 6.24 > 6, the evidence for the sublinear relationship against the linear relationship is strong^17^. We further studied how the total gene expression level scaled with chromosome size. Quantified by summing up the TPM (transcript per million) on each chromosome in a colon cancer cell line GP5d, we found that total gene expression also scaled sublinearly, with a scaling exponent of 0.65 (ΔBIC = 2.54, positive evidence, **Fig. 2b**).

On the basis of the same assumption, we would expect small TFs and large transcription complexes to differ in their scaling laws with respect to chromosome size. Indeed, our analysis of ChIP-seq data exhibited distinct differences for small TFs capable of binding to closed chromatin and large complexes limited by size exclusion (**Fig. 2c-e**). We plotted the count of binding sites of small TFs and large complexes on each chromosome in logarithmic scale. The binding sites of large transcription complexes, including Mediator (ΔBIC = 3.13, positive evidence, **Fig. 2c**) and RNA Polymerase II (ΔBIC = 3.40, positive evidence, **Fig. 2d**) scaled sublinearly, consistent with their localization with open chromatin. In contrast, small TFs that can bind to closed chromatin regions, such as HLF (**Fig. 2e**) and JUND (**Extended Data Fig. 1**), scaled quasi-linearly with chromosome size. Note that not all TFs displayed a quasilinear scaling behavior. Instead, TF scaling was a continuum, when we plotted the power law exponents (slopes of the regression lines) for different TFs (**Extended Data Fig. 2**). We attributed such distinction in the scaling behavior of TFs to their varied preference for open chromatin regions. While TFs like HLF and JUND have no preference for binding open regions over closed regions, other TFs may preferentially bind to open chromatin regions, even though their binding to chromosomal interiors are not constrained by their physical size. The diverse scaling behavior of small TFs and large complexes further stated the effect of size exclusion in the interaction between DNA and TFs. To further test our hypothesis of sublinear scaling, we also studied how average gene length on a chromosome scales with the chromosome size. Indeed, it followed a sublinear scaling relationship, with the power law exponent close to ⅓, the length-to-volume ratio (ΔBIC = 25.61, decisive evidence, **Extended Data Fig. 3**), suggesting that the average length of genes that a chromosome contains correlates with the diameter of the chromosomal territory.

In addition to the continuous scaling relationships modeled by power law, we also addressed 3D packing of the chromosomal territory from a discrete point of view. We estimated the fraction of nucleosomes that reside on the surface of chromosome territories by modeling it as a close-packing of equal spheres. We found that, for large chromosomes such as chr1 and chr2, the fraction of surface nucleosomes is around 5%. As the size of chromosomes decreases, the fraction of nucleosomes residing on the surface increases, up to 8.7% for chr21 (**Fig. 2f**). The more exposed nucleosomes on smaller chromosomes reinforced our assumption that smaller chromosomes are more accessible to the transcription machinery, thus they get higher odds of being transcribed per unit length.

We next asked what would happen if chromosome size was adequately small that each chromosome would be uniformly accessible? Driven by such a question, we carried out a similar scaling analysis for the yeast *Saccharomyces cerevisiae*, that lack chromosome territories and their chromosomes are more loosely arranged^18^. We approximated transcriptional activity with the count of Open Reading Frames (ORFs) on each chromosome, then plotted it versus chromosome size in log-log scale. Intriguingly, it turned out that gene expression in yeast strongly correlates with chromosome size following a linear scaling law (**Extended Data Fig. 4**). Taken together, how transcriptional activity scales with chromosome size largely depends on its 3D organization, and the presence or absence of chromosome territories in the species.

### Chromatin accessibility is modulated at different scales

The consistency of sublinear scaling law of transcriptional activity and binding of transcriptional machinery, as well as the anti-correlation between chromosome size and fraction of nucleosomes on the surface, supported the potential role played by size exclusion in controlling gene expression. It also implied that the accessible chromatin landscape would differ for small and large transcriptional complexes. We thus proceeded to develop a scalable probe to directly assess chromatin accessibility for molecules of different sizes. Specifically, we conjugated hyperactive Tn5 transposase with gold nanoparticles which consist of a gold core and a surface coating. The gold core defines the fundamental characteristics of a gold nanoparticle, whereas the protective coating can be modified for the interaction with biological probes and for stability. Gold nanoparticles are commercially available in various, well-defined sizes, and we used nanoparticles of 5 nm, 10 nm, and 20 nm in diameter in our studies. Transposase can be covalently attached to the surface of the ultra-stable gold nanoparticles through lysine residues, allowing us to make the scalable probes of chromatin accessibility. The conjugation between transposase and desired size of gold nanoparticles was followed by a standard ATAC-seq procedure^8^: tagmentation of the genome by assembled transposome, purification of tagmented DNA, PCR amplification, and Illumina sequencing.

We found that, at the chromosomal level (∼100 million base pairs), the overall open chromatin landscapes to small and large molecules were strikingly similar, except for Tn5 conjugated with 20 nm gold nanoparticles where no distinctive peaks were observed (**Fig. 3a**). However, as we zoomed into a finer locus level (∼10 kilo base pairs), distinct patterns of accessibility emerged. For example, at CTSD and SCN2B gene loci, the open regions to free Tn5 showed a clear enrichment around promoters, regions accessible to larger gold conjugates were dispersed along gene bodies, with reduced accessibility as molecule size increased.

**Fig. 3:**
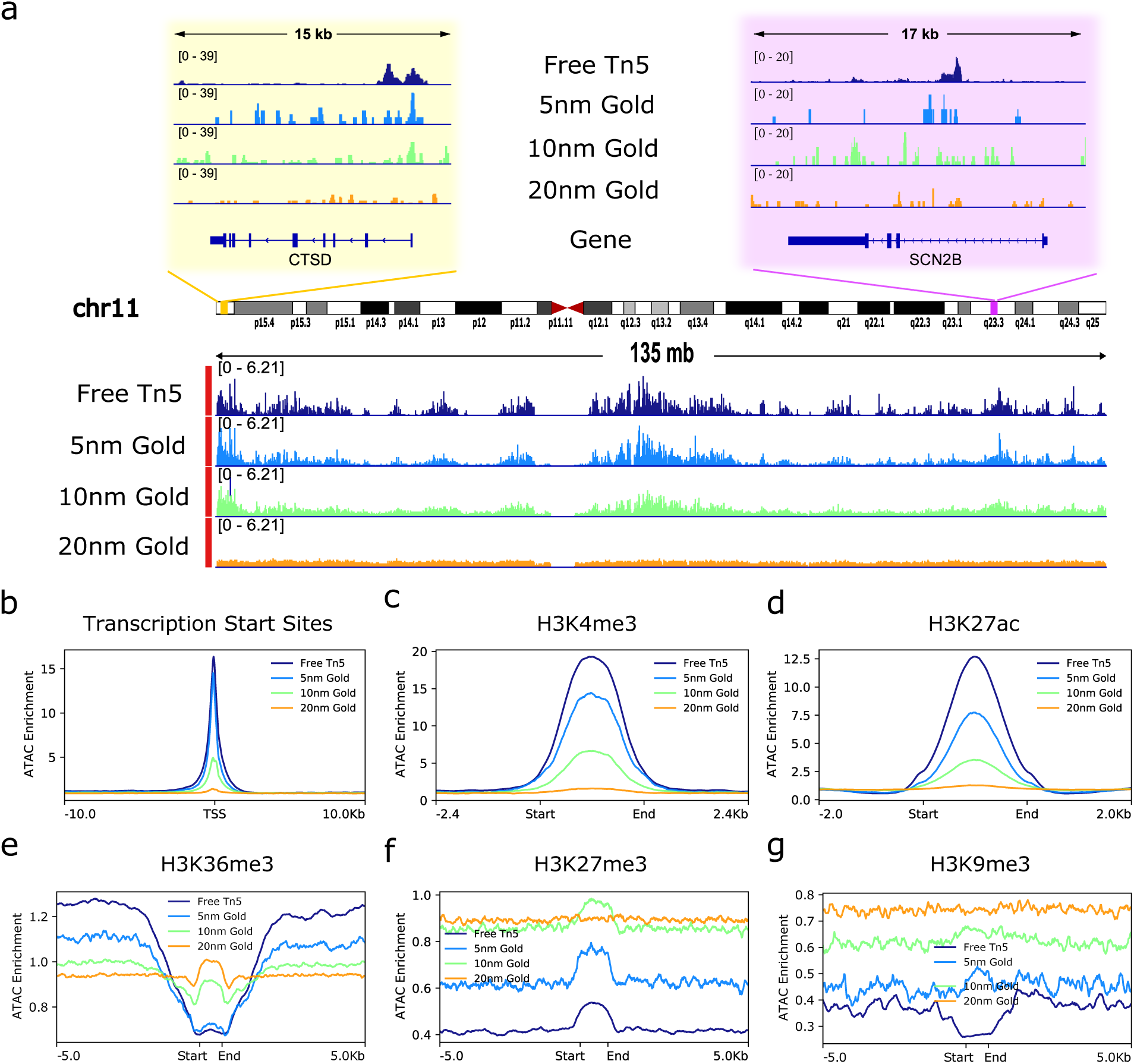
Chromatin accessibility at different scales, and ATAC-seq signal enrichment at different genomic features. **a,** Chromatin accessibility is controlled at two different scales - at chromosomal level, accessibility to small and large molecules is quite similar whereas at locus level, accessibility to small molecules was enriched at promoter regions and accessibility to large molecules was dispersed along gene bodies. **b-g,** ATAC-seq signal enrichment plots at Transcription Start Sites (**b**), H3K4me3 (**c**), H3K27ac (**d**), H3K36me3 (**e**), H3K27me3 (**f**), and H3K9me3 (**g**). Respective genomic features were first scaled to the same width (for **c-g**), then aligned by the reference point (for **b**) or by start and end points (for **c-g**). ATAC-seq reads mapped to regions within the genomic features were aggregated to display a global chromatin accessibility enrichment at various genomic features to molecules of different sizes.

### Subnuclear structures and chromatin accessibility

We next asked what key players underlie such distinct accessible patterns and may account for the chromatin accessibility to large molecules. As the cell nucleus is a complex and highly dynamic environment with many functionally specialized regions, we reason that subnuclear structures could play an important role in modulating chromatin accessibility. In particular, we were interested in the role of nuclear speckles and nuclear lamina. Nuclear speckles are granular-looking, membraneless condensates in the interchromatin space of the nucleus that contain high concentrations of RNA-processing factors and some TFs^19^. Recent studies have shown that the core of nuclear speckles is formed by two large, flexible proteins called SON and SRRM2^20^. On the functional side, nuclear speckles can enhance gene expression; may serve as storage and recycling sites for splicing factors; and might regulate the availability of splicing factors to control gene expression level^21,22^. In addition to its association with gene expression and splicing factors, we found that nuclear speckle associated domains (SPADs) can also facilitate chromatin accessibility (**Extended Data Fig. 5a**). We divided the ATAC-seq reads into two groups, the ones overlap with a SPAD and the ones do not. There turned out to be more reads in SPADs overlapping regions, irrespective of the probe size (Brunner-Munzel test with Bonferroni correction). Despite the positive correlation between the presence of SPADs and chromatin accessibility, we found that as we increased the probe size, the number of reads that were detected by the probe dramatically decreased (Kruskal-Wallis H test). We further investigated the effect of SPADs on chromatin accessibility by computing the global enrichment of ATAC-seq reads around SPADs. Specifically, we aligned all the SPADs in the whole genome and scaled them so that they have the same width. We then aggregated the ATAC-seq reads inside and flanking the SPADs (**Extended Data Fig. 5b**). We found that ATAC-seq signal enrichment peaked at the periphery of SPADs regardless of the probe size. While comparable ATAC-seq enrichment was found from free Tn5 and 5 nm gold conjugate probes, there was much reduced enrichment observed at the periphery of SPADs from 10 nm gold conjugate, with 20 nm gold conjugate showing unnoticeable peaks. This suggested that although SPADs may facilitate chromatin accessibility for small molecules, they have limited effect on large molecules.

Nuclear lamina is a structure on the inner surface of the cell nucleus and lamina-associated domains (LADs) are generally associated with gene repression and reduced chromatin accessibility^23^. We wondered whether the repressive effect would differ for molecules of different sizes. Therefore, we divided ATAC-seq reads into groups that reside within LADs and groups that reside outside LADs (**Extended Data Fig. 6a**). We found that the repressive effect of LADs held true for probes of all sizes (Brunner-Munzel test with Bonferroni correction). As expected, the read count decreased as the probe size increased (Kruskal-Wallis H test). We further studied the ATAC-seq enrichment around LADs and found clear reduced enrichment inside LADs from free Tn5 and 5 nm gold conjugate probes. The repressive effect was much alleviated for 10 nm gold conjugate, and surprisingly little for 20 nm gold conjugate (**Extended Data Fig. 6b**). It implied that LADs had selective repression over small and large molecules, and the commonly believed repressive effect of LADs may not account for the limited chromatin accessibility for large molecules.

### Genomic features and chromatin accessibility

In spite of the observed facilitative effect of SPADs and repressive effect of LADs, they didn’t decipher the chromatin accessibility to large molecules. Hence, we subsequently sought out the genomic features underlying the differentially accessible chromatin regions to molecules of different sizes. We first looked into ATAC-seq signal enrichment in TSSs (**Fig. 3b**), as transcription initiation closely relates to open chromatin. Specifically, we used TSSs as reference points and assessed the ATAC-seq enrichment 10 kb flanking the TSSs. We found that cells probed by free Tn5 showed a marginally higher ATAC-seq enrichment compared to cells probed by 5 nm gold conjugate. However, for larger molecules, chromatin accessibility dropped dramatically, reflected in the much lower peak for 10 nm gold conjugate and the hardly discernible peak for 20 nm gold conjugate. It suggested that TSSs are not always accessible to the larger molecules and transcription complexes, potentially due to a presence of existing large molecular complexes (such as paused polymerases) at these sites.

### Gene body exhibited higher chromatin accessibility for large molecules

Since histone marks are associated with distinct states of transcription, we subsequently studied chromatin accessibility at common transcriptional active histone marks, including H3K4me3 (**Fig. 3c**) and H3K27ac (**Fig. 3d**). We found that these transcriptional active regions exhibited decreased accessibility as the probe molecule size increased from native Tn5 to Tn5 conjugated with 10 nm gold; finally, for the 20 nm Tn5 gold conjugate, almost no ATAC-seq read enrichment was observed. On the contrary, the ATAC-seq enrichment in gene bodies, marked by histone modification H3K36me3, exhibited a completely different enrichment pattern (**Fig. 3e**). The gene body regions showed higher ATAC-seq enrichment for larger molecules, in contrast to the other active histone marks. This observation was also confirmed visually by inspecting ATAC-seq pattern at specific genes (example shown in **Fig. 3a**). The finding is unexpected, as gene bodies are not commonly associated with accessible chromatin. Our finding is, however, consistent with recent analysis of the spatial organization of transcribed eukaryotic genes, which suggest that long highly expressed genes form open-ended transcription loops, where polymerases move along the loops and carry nascent RNAs undergoing co-transcriptional splicing^15^. Such lateral loops emanating from the chromosome axis were observed before as lampbrush chromosomes^24^ and puffs of polytene chromosomes^25^. We reasoned that when gene bodies get transcribed, the moving Pol II with a cargo of nascent RNA could open up the original densely packed chromatin, leading to higher accessibility for large molecules. Indeed, TT-seq of nascent RNA verified that nascent RNA was enriched along gene bodies (**Extended Data Fig. 7**).

We next investigated the effect of heterochromatin on DNA accessibility. Analysis of ATAC-seq enrichment at H3K27me3 (**Fig. 3f**), PRC1 associated heterochromatin histone mark^26^, and H3K9me3 (**Fig. 3g**), HP1 associated heterochromatin histone mark^27^, revealed that both types of heterochromatin were generally not very accessible, in comparison with ATAC-seq enrichment scores of the other active histone marks. For free Tn5, 5 nm, and 10 nm gold conjugates, there was ATAC-seq enrichment within the position of the H3K27me3 mark regions compared to its flanking region. This enrichment was lost for the 20 nm Tn5 gold conjugate, suggesting that H3K27me3 modified chromatin is less accessible for molecules larger than 20 nm. For H3K9me3 containing regions, 5 nm gold conjugated Tn5 had ATAC-seq enrichment within H3K9me3 mark regions whereas chromatin accessibility of free Tn5 was noticeably repressed within H3K9me3 regions, probably due to a lower frequency of nucleosome-depleted regions at H3K9me3 heterochromatin. By contrast, the 10 nm and 20 nm conjugated Tn5 showed little to no enrichment, suggesting that H3K9me3 heterochromatin is inaccessible to large molecules.

## Discussion

In this work, we have developed scalable probes to measure chromatin accessibility to molecules of different sizes. To increase the size of molecules, we conjugated the Tn5 transposase with different sizes of gold nanoparticles, which are highly uniform in size, and biochemically inert. We found that the accessible regions for small molecules were primarily enriched at promoter regions, a classical open chromatin region, whereas accessible regions for large molecules were dispersed along gene bodies. We observed a close to ⅔ sublinear scaling of transcriptional activity, and of large transcriptional complexes binding to DNA, and close to ⅓ sublinear scaling of gene length. These results suggest that transcriptional activity and gene length in a chromosome is limited by the size of the chromosome territory.

Previous studies have analyzed diffusion of macromolecules of different sizes in the nucleus^28^, using GFP multimers and dextrans as probes. The radius of the probes ranged from ∼3 nm (GFP monomer) to ∼45 nm (500 kDa dextran). In these studies, the authors observed reduced concentration of the probes in heterochromatin compared to euchromatin due to size exclusion, but also observed that even the densest nuclear compartments are highly permeable, and readily accessible to diffusing proteins. The authors reported the estimated sizes of their larger dextran probes based on radius of gyration, which overestimates the hydrodynamic size of large molecules. By converting to Stokes-Einstein radius, which is related to size exclusion, the size range of the molecules used in the studies were similar to those used here, making the earlier observations broadly consistent with our findings. Furthermore, studies have used electron microscopy tomography to visualize 3D chromatin structure in interphase nuclei^29^, finding that the percentage of chromatin volume in nuclear volume exhibit a normal distribution ranging from 12% to 52%, with heterochromatin domains at the nuclear envelope showing higher chromatin volume concentration, from 37% to 52%. These observations are also broadly consistent with our work.

Our findings, together with data from the recent studies of the spatial organization of transcribed eukaryotic genes^15^, and microphase separation of RNA and DNA in euchromatin^30^, suggest a working model for the role of size exclusion in chromosome structure and gene expression (**Fig. 4**). In the model, chromatin prefers to associate with chromatin, but not with disordered polymers such as RNA. This model predicts two distinct nuclear domains: the chromatin phase containing relatively densely packed DNA, and the RNA phase containing nascent RNA and other large macromolecules that are largely excluded from the interior of the chromosome territories. Genes undergoing transcription, however, are not purely DNA, nor purely RNA. Rather, they exist as a hybrid structure that naturally associates with the interphase between RNA and chromatin. This model also fits in the observed accessibility of gene bodies to large molecules, resulting from large nascent RNA molecules needing to exit their gene bodies. For example, the hydrodynamic radius of an 8.9 kb single-stranded RNA is ∼33 nm^31^, similar to that of Mediator/RNA PolII complex. Transcriptional activity in such a system would result in nascent RNA accumulation within the RNA phase, where large molecules such as the nascent RNA and Mediator/RNA PolII complex can diffuse relatively freely. This accumulation could also cause a stretching and unwinding effect on the original densely packed chromatin. Collectively, this intricate interplay would then culminate in the positioning of transcribed genes at the surfaces of chromosomes.

**Fig. 4:**
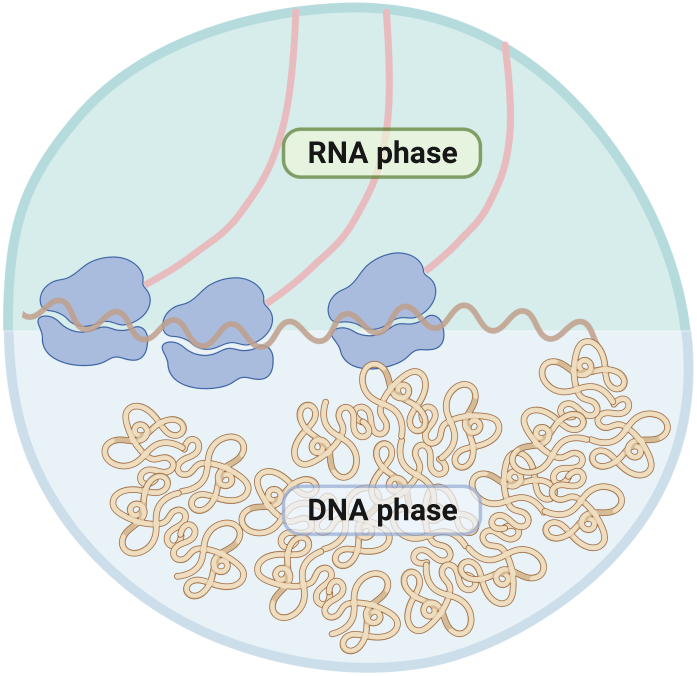
A working model for gene bodies to be accessible to large molecules during transcription. After transcription onset, RNA polymerase II moves along the gene axis and carries cargos of nascent RNA together with RNA binding proteins undergoing co-transcriptional splicing. The bulky nascent ribonucleoproteins on one hand increased the stiffness of the genes, on the other hand segregated from domains of transcriptionally inactive chromatin, establishing a distinct transcriptionally active domain which is enriched with RNA and RNA-binding proteins. Transcription continues at the interface of chromatin and RNA domains, providing increased accessibility for large molecules.

Inside cells, the surface of chromosome territory is uneven and textured, and dynamic to the extent that in addition to surface nucleosomes, many nucleosomes located close to the surface are also accessible to large molecules. Using a smooth spherical surface to model chromosomes is thus oversimplified. However, permeability of the surface, or mottles and dapples in it will not affect the surface to volume scaling of transcriptional activity, provided that the chromosomes are sufficiently large to contain an interior that is inaccessible to large molecules.

Our results also complement earlier functional genomic studies of nuclear structure. For example, a recently developed method by sequencing gradually digested chromatin along the nuclear radius has unveiled the radial organization of chromatin in the cell nucleus with improved resolution^32^. While it shows that chromatin accessibility increases globally toward the nuclear interior when compared to the nuclear periphery, this does not conflict with our findings, as we focus on the surface of chromosome territories, and the nuclear interior is likely where surfaces of different chromosomes touch and interact with one another. Furthermore, studies of chromatin tracing and live-cell imaging have shown that genes are repositioned toward the centroid of the surrounding chromatin upon activation^33^. It is worth noting that the centroid was calculated locally for genomic windows spanning several megabases, distinguishing it from the centroid of intact chromosomes. While this suggested a spatial coupling between transcribed genes and the formation of transcription factories, whether such repositioning extends to the entire chromosomal level remains to be investigated.

Our work is also relevant to the mechanisms that control gene expression. Earlier, Maeshima et al. noted the difference in sizes of macromolecules involved in transcription^7^, and proposed a transcriptional activation model where enhancer acts as a buoy, interacting with gene promoters and causing the latter to float up to the surface from the inner core of the condensed domains^34^. This suggests that large chromosome size would enable a new level of regulation of gene expression. Here we show that size exclusion effect operates at the chromosome level, separately from the local effect of nucleosome occupancy on binding of TFs to DNA. Local nucleosome clustering, as observed for example in heterochromatin, can also selectively affect diffusion of large molecules due to steric hindrance. However, the scaling laws we have observed in relation to varying chromosome sizes strongly suggests a regulation mechanism also operates at the whole chromosome level.

The recently identified chromatin context-dependent enhancers^35^ and “facilitator” enhancer elements^36^ have provided suggestive evidence for the model that enhancers can function by regulating access of large molecules, and by bringing a locus to the surface of chromosomes. In particular, absence of the facilitator elements severely reduced binding of transcriptional coactivators including Med1 and Brd4 (bromodomain-containing protein 4), but not binding of TFs, suggesting that the facilitator enhancers are chromatin-dependent elements that facilitate gene expression via chromatin modification or structural changes in 3D organization of chromatin.

The chromatin-dependent distal regulatory elements would not be necessary for organisms such as the yeast *S. cerevisiae* that have small chromosomes. However, the elements would enable chromosome size to be increased dramatically, as the interior would not be always inactive, but could be activated by a regulatory mechanism. Consistent with this hypothesis, we found here that in contrast to human chromosomes, yeast chromosomes do not display sublinear scaling. This observation potentially explains why yeast does not have enhancers, despite having all the molecular machinery, such as Mediator complex, that are commonly associated with enhancer activity.

Taken together, our work uncovered a surprising link between chromosome size and gene expression. The identified scaling laws have wide implications to chromosome structure, gene expression and evolution of complex organisms. Further mechanistic analysis of the scaling laws by structural and imaging technologies will undoubtedly lead to a more comprehensive understanding of gene expression, and potentially pinpoint the mechanism of action of a critical gene regulatory element, the enhancer.

## Figures

**Extended Data Fig. 1:**
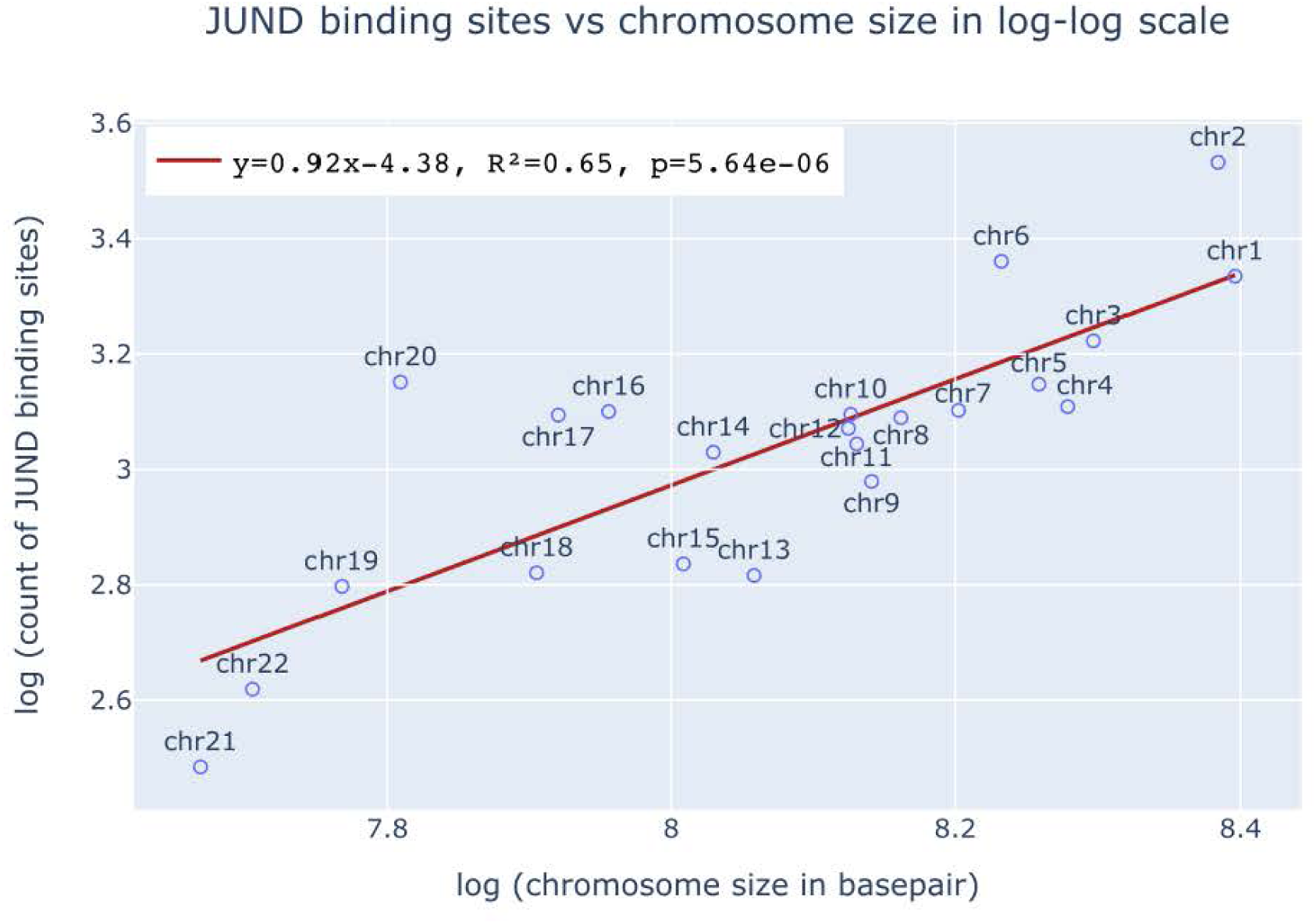
The count of JUND binding sites scales quasi-linearly to chromosome size with an exponent close to 1. The count of JUND binding sites was determined from ChIP-seq data. Red solid line denotes the regression line between JUND binding sites and chromosome size in logarithmic scale.

**Extended Data Fig. 2:**
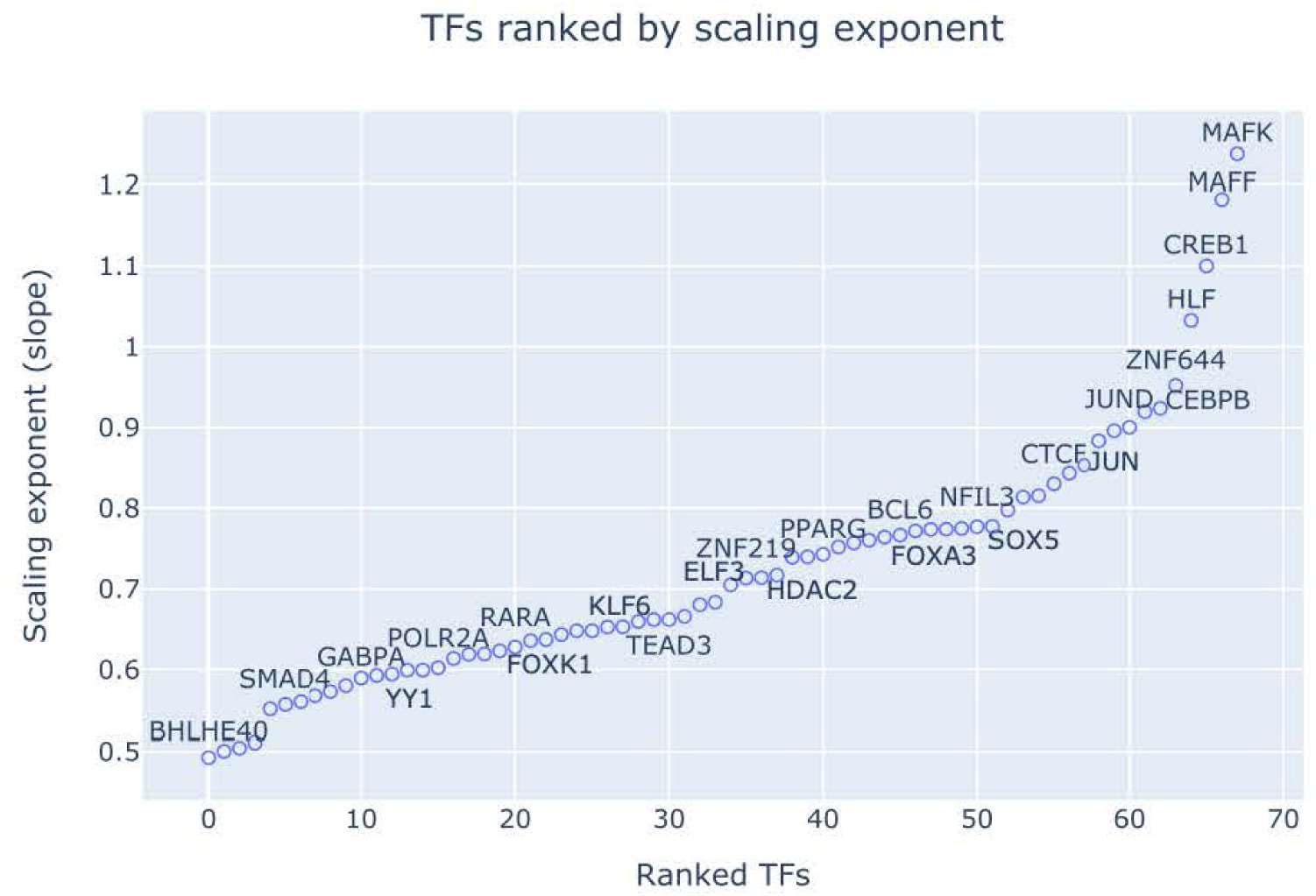
Scaling exponents of various TFs. The counts of binding sites on each chromosome were determined for 128 TFs (including large complexes like POLR2A) using ChIP-seq data in the HepG2 cell line. Non-constrained regression was performed for each TF individually in logarithmic scale and the TFs were sorted by the ascending order of slopes of the regression line. TFs with very low R^2^ values (R^2^ <= 0.2) were filtered out (68 TFs left).

**Extended Data Fig. 3:**
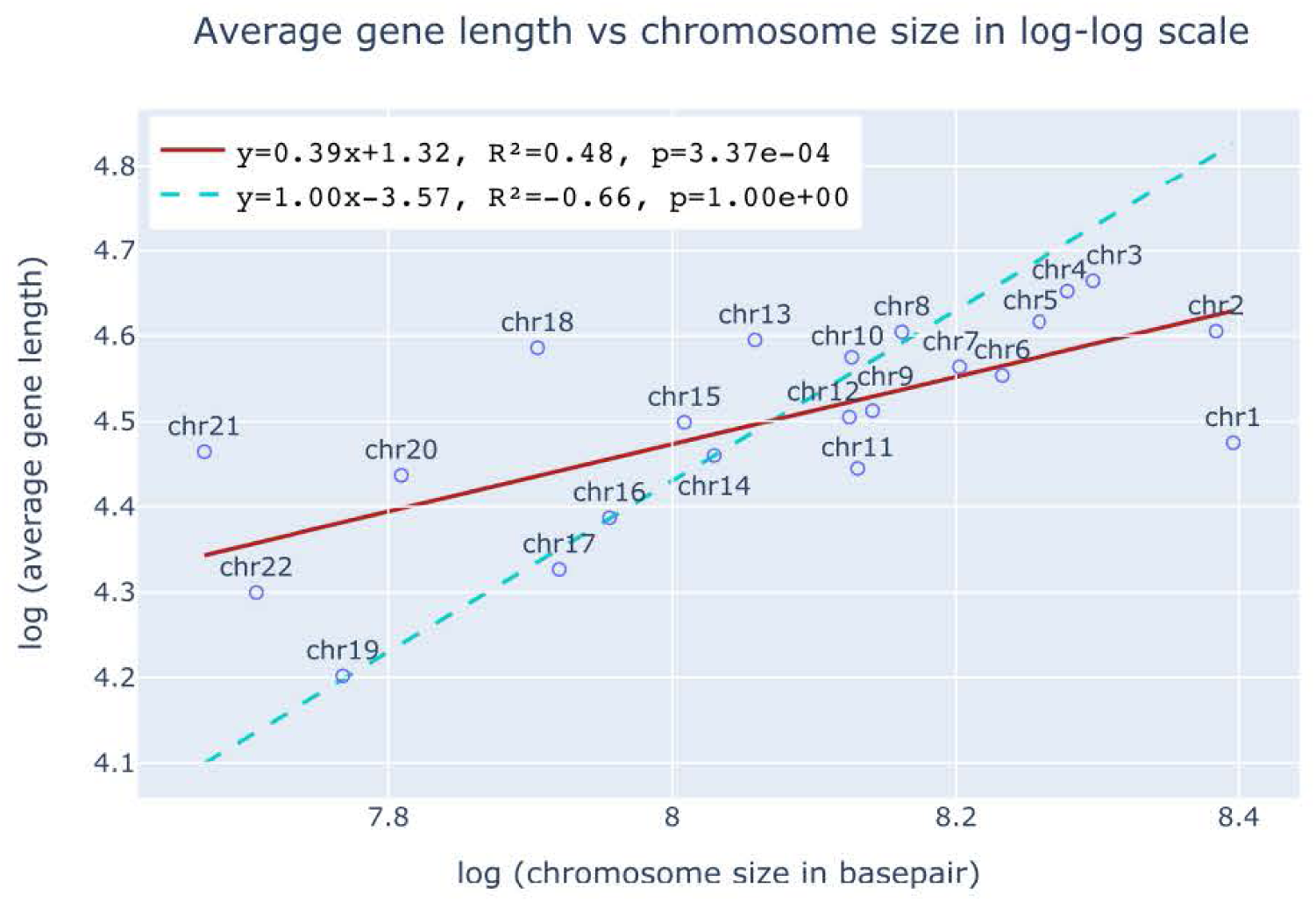
Average gene length scales sublinearly to chromosome size with an exponent close to ⅓. Red solid line denotes the regression line between average gene length on each chromosome and chromosome size in logarithmic scale. Cyan dashed line denotes the constrained linear regression line (slope is constrained to be 1 while intercept is optimized to best fit the data).

**Extended Data Fig. 4:**
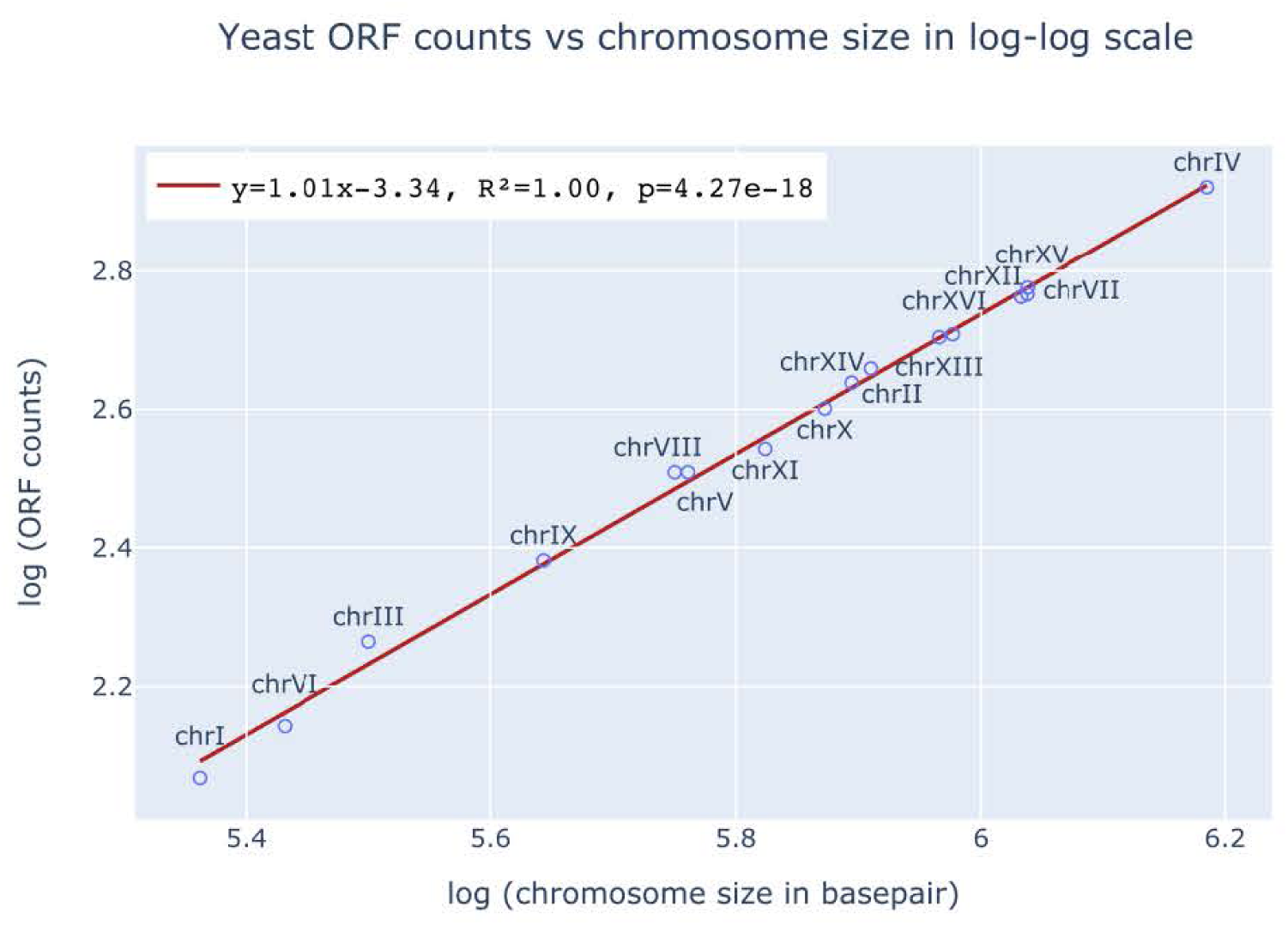
The count of ORF in yeast scales linearly to chromosome size with an exponent close to 1. The count of Open Reading Frame (ORF) on each chromosome in yeast was used as a proxy of transcriptional activity. The slope of the optimal non-constrained regression line was quite close to 1, indicating a linear scaling relationship between ORF count and chromosome size in yeast.

**Extended Data Fig. 5:**
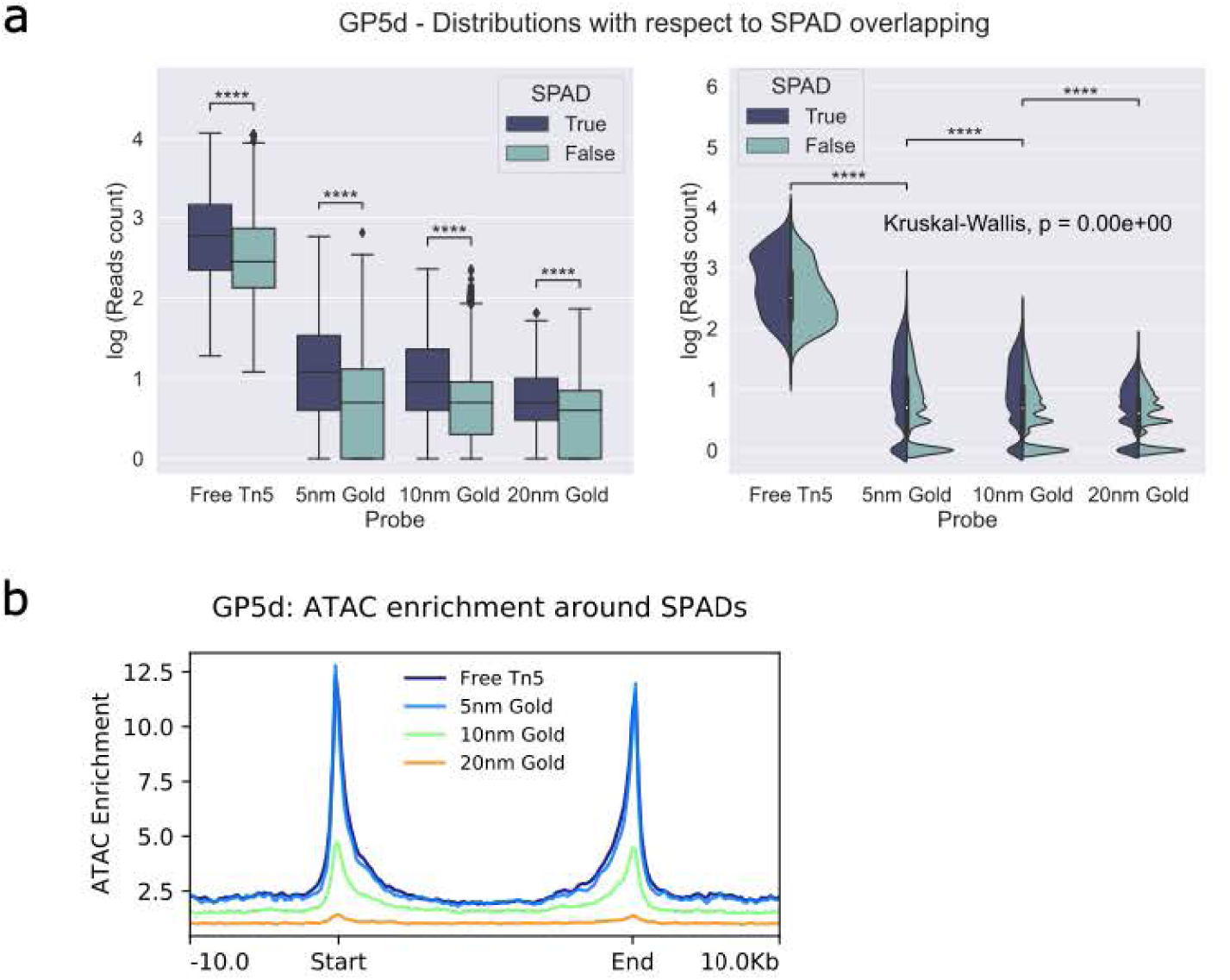
Effect of Nuclear Speckle Associated Domains (SPADs) on chromatin accessibility to molecules of different sizes. **a,** Distribution of log transformed reads count with respect to SPAD overlapping in GP5d cell line. Boxplot on the left shows pairwise comparison for loci that overlap with SPADs and those that do not. P values were calculated based on Brunner-Munzel test with Bonferroni correction. p=0.000e+00 for free Tn5, 5 nm, and 10 nm gold conjugates; p=1.172e-266 for 20 nm gold conjugate. Violin plot on the right shows distribution of log transformed reads count that was probed by molecules of different sizes. Kruskal-Wallis H test was performed for all the 4 groups and global p=0.00e+00. Brunner-Munzel test with Bonferroni correction was then performed for pairwise comparison between adjacent groups individually. p=0.000e+00 for Free Tn5 versus 5 nm gold conjugate, and for 10 nm gold conjugate versus 20 nm gold conjugate; p=5.339e-130 for 5 nm gold conjugate versus 10 nm gold conjugate. **b,** ATAC-seq signal enrichment plot around SPADs. SPADs were first scaled to the same width, then aligned by start and end points. ATAC-seq reads mapped to regions within SPADs were aggregated to display a global chromatin accessibility enrichment within/flanking SPADs to molecules of different sizes.

**Extended Data Fig. 6:**
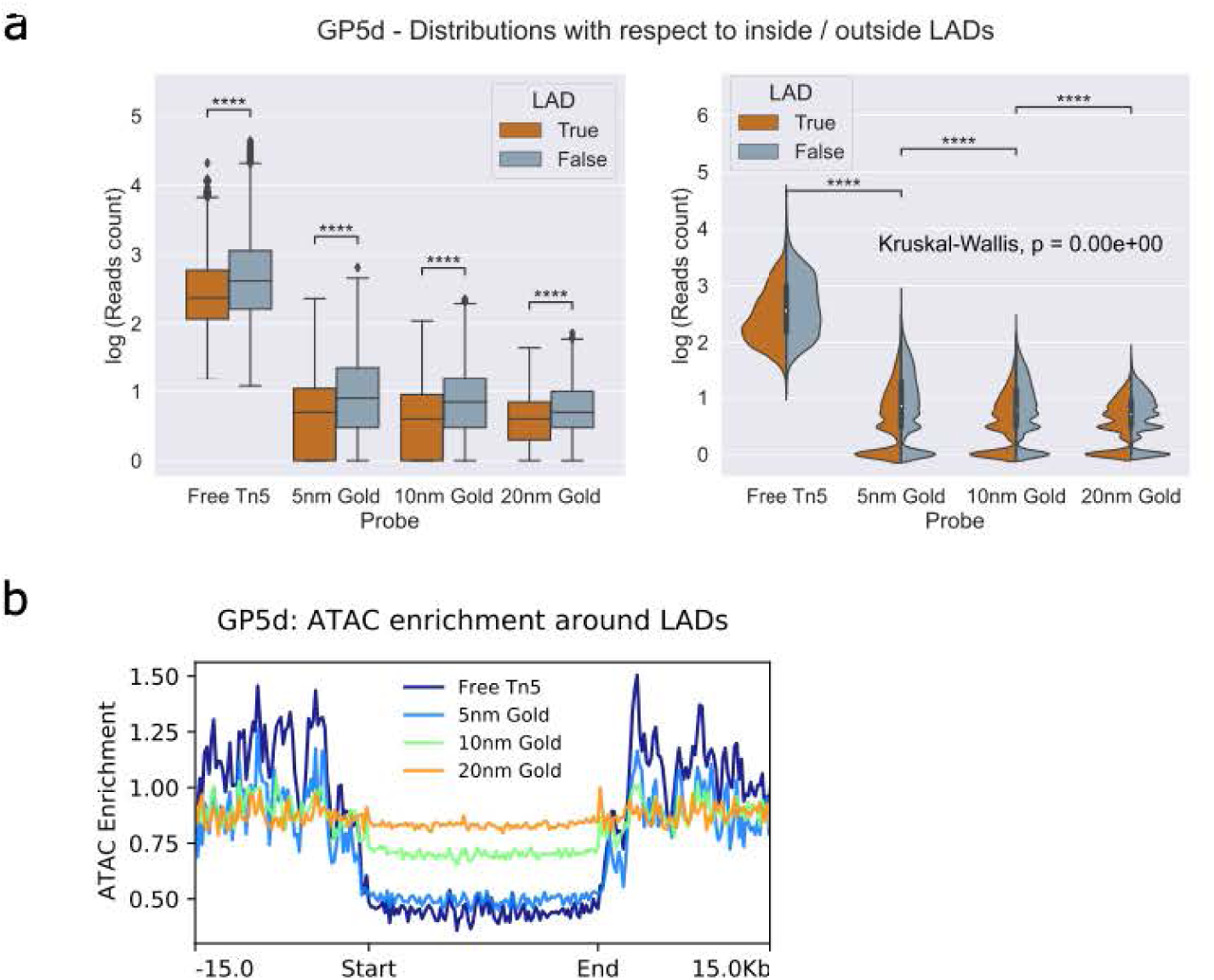
Effect of Lamina Associated Domains (LADs) on chromatin accessibility to molecules of different sizes. **a,** Distribution of log transformed reads count with respect to inside/outside LADs in GP5d cell line. Boxplot on the left shows pairwise comparison for loci that are inside and that are outside LADs. P values were calculated based on Brunner-Munzel test with Bonferroni correction. p=0.000e+00 for all the probes. Violin plot on the right shows distribution of log transformed reads count that was probed by molecules of different sizes. Kruskal-Wallis H test was performed for all the 4 groups and global p=0.00e+00. Brunner-Munzel test with Bonferroni correction was then performed for pairwise comparison between adjacent groups individually. p=0.000e+00 for Free Tn5 versus 5 nm gold conjugate, and for 10 nm gold conjugate versus 20 nm gold conjugate; p=1.384e-123 for 5 nm gold conjugate versus 10 nm gold conjugate. **b,** ATAC-seq signal enrichment plot around LADs. LADs were first scaled to the same width, then aligned by start and end points. ATAC-seq reads mapped to regions within LADs were aggregated to display a global chromatin accessibility enrichment within/flanking LADs to molecules of different sizes.

**Extended Data Fig. 7:**
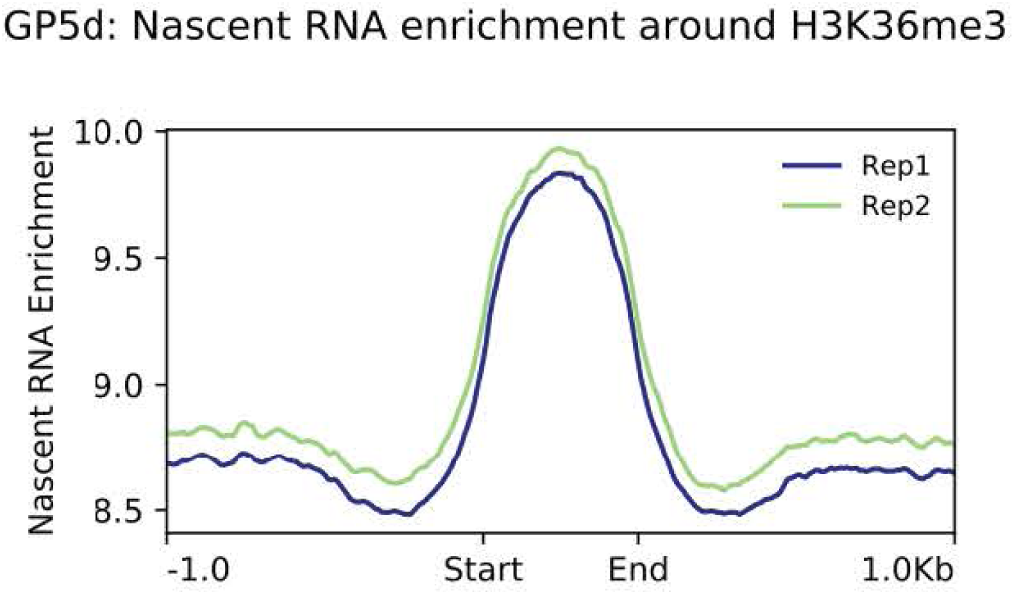
Nascent RNA enrichment plot at H3K36me3. Nascent RNA enrichment was computed by counting TT-seq reads in GP5d cell line (two biological replicates). H3K36me3 marked sites were first scaled to the same width, then aligned by start and end points. TT-seq reads mapped to regions with H3K36me3 modification were aggregated to display a global nascent RNA enrichment within/flanking H3K36me3 modified loci.

## Methods

### Cell culture

The colon cancer cell line GP5d (Sigma-Aldrich, catalog no. 95090715) was cultured in DMEM, supplemented with 10% fetal bovine serum, 2 nM L-glutamine and 1% penicillin-streptomycin. Cells were examined for mycoplasma contamination regularly. Briefly, cells were plated on gelatin-coated coverslips in 12-well plates, fixed with 4% PFA for 10 mins and stained with Hoechst, then observed under UV channel of the microscope.

### Gold conjugation

Assembled Tn5 Transposome (Active Motif, 53150) underwent buffer exchange with spin desalting columns (Thermo Fisher, 89882) first, to get rid of primary amines that may interfere with the covalent reaction. The Transposome was loaded in amine-free buffer, 10 mM HEPES, supplemented with Protease inhibitor (Sigma-Aldrich, 11873580001). Gold conjugation was performed with 5 nm (Sigma-Aldrich, 900458), 10 nm (abcam, ab201808), and 20 nm (abcam, ab188215) nanoparticles. The Tn5 Transposome was combined with gold reaction buffer in a proportion specific to the conjugation kit used, the ligand solution was subsequently transferred to a vial containing gold nanoparticles and mixed well. The conjugation reaction was incubated at room temperature for 0.5-1 h depending on the kit used. A quencher solution was then added to stop the reaction. As a conjugate 100% free from unbound Transposome is desired, the conjugate was then washed with 1:10 diluted quencher and centrifuged according to the kit manual. The supernatant was carefully removed and the gold conjugate was stored at 4℃ until use.

### ATAC-seq and sequencing library preparation

ATAC-seq was performed according to the kit manual (Active Motif, 53150). Briefly, cells were counted with Countess 3 Automated Cell Counter (Thermo Fisher) and around 100,000 cells were allotted into a fresh 1.5 mL tube and centrifuged at 500 g for 10 min at 4℃. Supernatant was removed and ice-cold PBS was added, followed by a spin-down at 500 g for 5 min at 4℃. Cell pellet was then resuspended in 100 µL ice-cold ATAC lysis buffer and spun down at 500 g for 10 min at 4℃. Supernatant was removed and tagmentation mix solution was added. The tagmentation reaction was incubated in a thermomixer at 800 rpm for 1h at 37℃. DNA purification binding buffer was then added, and DNA purification and elution was performed with DNA purification columns. PCR amplification of the purified DNA was then performed, followed by size selection with SPRI beads. The size distribution was then assessed with TapeStation (Agilent) and the DNA was sequenced with Illumina NextSeq500 (paired-end sequencing, 300-cycles Mid Output Kit).

### ATAC-seq data analysis and signal enrichment plot

The paired-end reads (fastq format) were mapped to hg38 using Bowtie2 (v2.3.1)^37^ performing local alignment with parameters –very sensitive. Over ∼7.9 million mapped reads were generated in each sequencing library and used for downstream data mining. Samtools (v1.7)^38^ were used to sort and compress the resultant SAM files into BAM files. Unmapped reads, duplicated reads, or reads of poor mapping quality were filtered out. Coverage tracks (bigWig format) were generated using bamCovergae (v3.5.2)^39^ and ENCODE blacklisted regions were removed. Coverage tracks were binned (binSize=10 bases) and normalized using RPGC (reads per genomic content), then inspected with Integrative Genomics Viewer (IGV, v2.11.1) or used to calculate enrichment scores per genomic region. RPGC (per bin) = number of reads per bin / scaling factor for 1x average coverage, where scaling factor is determined from the sequencing depth: (total number of mapped reads * fragment length) / effective genome size.

Broad-peak calling was performed with MACS3 (v3.0.0a6), and blacklisted regions were removed from peak files with bedtools (v2.20.1)^40^. Peaks called from standard ATAC-seq procedure (samples probed with free Tn5) were used as consensus peak list. Bedtools bamtobed was used for format conversion, and number of unique-mapped and properly paired reads mapped to each peak for each individual sample was quantified for downstream statistical analyses.

deepTools (v3.5.1)^39^ computeMatrix and plotProfile were used to make the ATAC-seq signal enrichment plots. Briefly, computeMatrix goes over a set of genomic features (e.g., TSSs, histone marks, SPADs/LADs) and coverage tracks (bigWig files prepared as described earlier). Genomic features (regions we are interested in) were first scaled to the same width which was then divided by the bin size to get the number of bins in each region that we need to compute enrichment values for. bigWig files were then used to extract all of the per-base values in the region. The per-base values were divided into bins, and the average in each bin in each region was recorded in a matrix. As the regions were all scaled and aligned with the same number of bins, plotProfile then calculated the average of each bin across regions and plotted. Samples probed with molecules of different sizes were superimposed to be displayed together.

### Computational analysis of chromatin

TSS data was obtained from Ensembl biomart (Ensembl Genes 110, Human genes GRCh38.p14, retrieved in March 2021), then we counted the number of TSSs on each chromosome. Gene start and end coordinates were obtained from Ensembl biomart, then we subtracted start from end coordinates to get gene length. RNA-seq data of GP5d was obtained from a previous study^41^, and Transcript Per Million (TPM) was used to quantify gene expression level. We then summed up TPMs of all the protein coding genes on each chromosome. ChIP-seq data of the TFs was obtained from a previous study^42^. Peak enrichment was computed using Phantompeakqualtools^43^, implementing SPP peak calling algorithm to generate narrow peaks. Irreproducible Discovery Rate (IDR)^44^ was run on replicates peak file and pooled peaks, using IDR-2.0.2 tool, to generate peaks passing IDR cutoff 0.02 (soft-idr-threshold) followed by filtering out ENCODE blacklisted regions. We then counted binding sites of individual TF on each chromosome. Yeast ORF data was obtained from *Saccharomyces* Genome Database, then we counted the number of ORFs on each chromosome.

To approximate the fraction of nucleosomes that reside on the surface of the chromosome territory, we modeled it as a close-packing of equal spheres problem. Each chromosome size in base pairs was first divided by 200 to approximate the number of nucleosomes in that chromosome. Suppose a chromosome territory is stuffed with DNA wrapping around *N* nucleosomes, we have *N* = 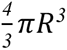 ∗ *d*_3_, where *R* is the radius of the chromosome territory in nucleosome width, and *d_3_* is the packing density of the three-dimensional densest regular packing arrangement, which is around 0.74. Suppose there are *n* nucleosomes reside on the surface of the territory, we have *n* = *4*π*R*^2^ ∗ *d_2_*, where *d_2_* is the packing density of the two-dimensional densest packing arrangement, which is around 0.91. We can then approximate the fraction of nucleosomes on the surface by taking 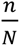.

### Statistical analysis of the scaling laws relative to chromosome size

All the scaling analysis was first modeled by non-constrained linear regression in log-log scale, and the coefficients (the slope and the intercept) of the regression line were determined by ordinary least squares method. F-statistic was used to test the significance of the coefficients of the indicator variable, and p-value indicated the significance of the slope.

As the optimal non-constrained regression for TSS count, total gene expression, MED1 binding sites, POLR2A binding sites, and average gene length implied a sublinear scaling relationship with chromosome size, the scaling of these variables was subsequently modeled by constrained regression, where the slope was fixed to 1 with the intercept being optimized to best fit the data. Bayesian information criterion (BIC) was then used to compare the linear scaling model and the sublinear scaling model. BIC is defined as *BIC* = *k ln* (*n*) − *2 ln* (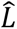). where *k* is the number of coefficients in each model, *n* is the number of data points, and 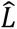 is the maximized value of the likelihood function of each model. ΔBIC between 6 and 10 means the evidence for the alternative model (sublinear scaling) against the null model (linear scaling) is strong, whereas a ΔBIC greater than 10 implies the evidence favoring the alternative model is decisive, and a ΔBIC smaller than 6 suggests a positive evidence for the alternative model^17^.

### Code availability

All custom code that supports the findings of this study is available at https://github.com/JiayinHong/Scaling-laws-of-transcription.

## Acknowledgments

We thank Ken Jones for assistance with cell culture, Teemu Kivioja, Kimmo Palin, Tuomo Hartonen, Ekaterina Morgunova, Shreya Saha, Ville Taipale, Aleksandra Jartseva, and Rui Guan for their input and discussions. We also thank Helen Mott and Xiaodan Liu for their help with protein expression, and Inderpreet Kaur Sur and Connor Rogerson for feedback on the manuscript. This work was supported by Cancer Research UK.

## Author contributions

J.H. and J.T conceptualized the study. J.H. performed experiments with assistance from A.D.C and D.S. J.H. performed analyses. J.H., J.T. and E.L. interpreted the data. J.H. and J.T. wrote and edited the manuscript, with feedback from E.L.

## Competing interests

The authors declare no competing interests.

## Materials & Correspondence

Correspondence to Jussi Taipale.

## Data availability

Source data are provided with this paper.

## Notes

### Competing Interest Statement

The authors have declared no competing interest.

### Summary of Updates

Discussion revised.

